# A Drosophila Tonic Motor Neuron Reinnervates Ectopic Muscles Fully Deprived of Native Tonic and Phasic Inputs

**DOI:** 10.1101/2025.07.25.666868

**Authors:** Lizzy Olsen, Parinita Mitchelle Mandhyan, Sara Arredondo, Komal Kaur, Ankura Sitaula, Aref Zarin

## Abstract

Motor neurons (MNs) form precise neuromuscular junctions (NMJs) during development, but the extent to which individual MNs can reinnervate fully denervated muscles in vivo remains poorly understood. In the Drosophila larva, each muscle is co-innervated by a tonic and a phasic glutamatergic MN. Here, we show that the tonic MN1 undergoes robust heterosynaptic sprouting and forms ectopic NMJs when neighboring muscles are deprived of both their native tonic and phasic inputs. This structural plasticity is not induced by silencing, but instead requires the physical ablation of adjacent MNs. Live imaging of the same MN1 axons in individual animals reveals that sprouting initiates early and expands progressively across larval stages. In contrast, phasic MNs show minimal remodeling, indicating that tonic MNs possess a greater intrinsic capacity for neuroplasticity. Notably, as MN1 establishes new synapses on targets it does not normally innervate, it redistributes pre-synaptic territory across both native and ectopic muscles. These findings identify a subtype-specific, injury-induced rewiring program in an intact motor circuit.

## Introduction

Spinal cord injury, stroke, and neurodegenerative disease disrupt neural circuits, impairing motor and cognitive function. Functional recovery may depend on heterosynaptic structural plasticity (HTSP), in which intact neurons sprout new branches to reinnervate denervated targets (1–3). Therapeutic strategies aimed at enhancing HTSP hold promise, but their success hinges on a deeper understanding of the intrinsic and extrinsic factors that promote or constrain neuronal sprouting (4). Neurons differ in properties such as firing dynamics (e.g., tonic versus phasic), neurotransmitter identity, and synaptic architecture (e.g., large versus small boutons), but it remains unclear whether all neuron types are equally capable of structural remodeling after injury. This gap limits our ability to predict which neurons can compensate for lost inputs—and under what conditions. A central question, therefore, is how physiologically distinct neurons respond to the dysfunction or loss of their synaptic partners. Here, we examine the capacity for post-injury axonal sprouting and neuromuscular junction (NMJ) remodeling in physiologically distinct tonic and phasic motor neurons in Drosophila larvae.

The *Drosophila* larval neuromuscular system provides a genetically accessible and anatomically tractable model for investigating neuron subtype-specific injury responses. Larval muscles are innervated by two classes of glutamatergic excitatory motor neurons (MNs): tonic-firing type Ib MNs, which form large boutons and typically innervate a single muscle, and phasic-firing type Is MNs, which form smaller boutons and innervate multiple muscles (5–12). These MNs converge on shared target muscles, creating a synaptic motif in which one tonic and one phasic MN co-innervate the same postsynaptic muscle fiber. This configuration provides a unique opportunity to directly compare how distinct MN subtypes respond to the same insult. Notably, convergence of tonic and phasic inputs onto a common postsynaptic cell is an evolutionarily conserved circuit motif observed across both vertebrate and invertebrate systems (13–16). Due to its simplicity and powerful genetic toolkit, the *Drosophila* NMJ has long served as a leading model for dissecting conserved mechanisms of synaptic development and plasticity at single-cell resolution (5, 6, 8, 10, 17–29).

Previous studies have shown that ablating—but not silencing—the phasic MN RP2 induces structural expansion of the NMJ formed by its co-innervating tonic partner MN1 on Muscle 1, while loss of MN1 has no effect on RP2 (7–10). These findings suggest that tonic MNs are more structurally adaptive than phasic MNs. However, since only one MN type was disrupted in those studies, the muscles retained partial excitatory input. Thus, whether complete denervation prompts more robust or qualitatively different plastic responses—and which neurons are capable of compensating—remains unknown.

To address this, we developed a genetic paradigm to simultaneously ablate both tonic and phasic MNs that co-innervate dorsal longitudinal muscles, creating a fully denervated group of target muscles. We found that the surviving tonic MN1, which normally innervates Muscle 1, exhibited robust axonal sprouting and formed ectopic NMJs on denervated muscles it does not typically target. In contrast, the phasic MN RP2 did not sprout following ablation of its tonic partners, and MN1 failed to sprout when neighboring MNs were chronically or acutely silenced rather than ablated. These results demonstrate that injury-induced HTSP is neuron-subtype specific and requires complete physical loss of input—not merely loss of activity—to initiate structural remodeling.

We further show that the ectopic NMJs formed by MN1 are functional: they restore calcium transients in previously denervated muscles and partially rescue crawling performance. Using live motor axon imaging, behavioral assays, and temporally controlled genetic ablation, we mapped the dynamics of MN1 sprouting across larval stages. Sprouting was initiated shortly after larval hatching and continued throughout larval development, revealing a broader plasticity window than previously appreciated.

Together, our findings identify a MN subtype-specific and injury-dependent form of HTSP that enables functional circuit compensation. This work uncovers key cellular and temporal principles governing adaptive circuit remodeling and provides a foundation for understanding how defined neurons sense local denervation and initiate compensatory growth programs.

## Results

### Tonic MN1 Sprouts and Forms Ectopic NMJs Following Complete Denervation of Neighboring Muscles

The *Drosophila* larval body comprises three thoracic segments (T1–T3) and nine abdominal segments (A1–A9). The central nervous system (CNS) consists of two brain lobes and a segmented ventral nerve cord (VNC), which functions analogously to the vertebrate spinal cord. Each VNC segment controls the muscles of its corresponding body segment (**Fig. 1A**). In each abdominal segment, approximately 30 bilaterally symmetrical muscle pairs are innervated by 28 pairs of tonic and two pairs of phasic glutamatergic excitatory motor neurons (MNs) projecting from the segmental VNC (5, 30–36) (**Fig. 1A**).

**Figure 1.**
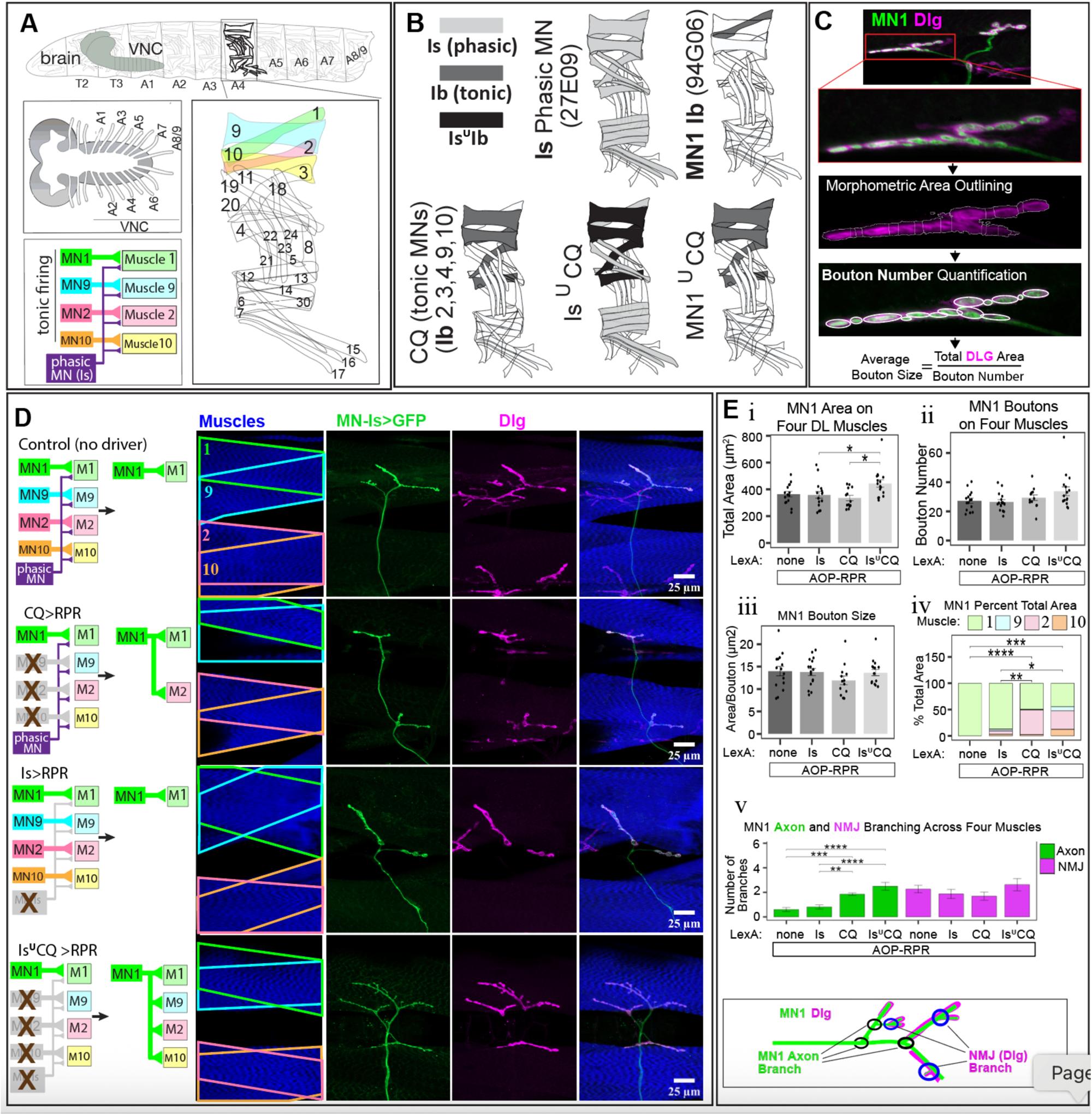
The tonic MN1 axon sprouts and forms ectopic NMJs on fully denervated dorsal-longitudinal (DL) muscles. (A) Schematic diagrams of the Drosophila larval nervous system and body wall muscles. This study focuses on four DL muscles (1, 2, 9, and 10), each receiving phasic input from a common phasic MN-Is and tonic input from distinct tonic Ib MNs (MN1, MN2, MN9, and MN10). (B) Target muscles of the Gal4 and LexA MN drivers used in this study. The union symbol (∪) indicates fly lines carrying two drivers simultaneously. For example, in *Is ∪ CQ* larvae, both tonic and phasic MNs innervating muscles 2, 3, 4, 9, and 10 are targeted. (C) Workflow illustrating the NMJ analysis pipeline [adapted from (41, 42)]. (D) Representative NMJ immunostaining visualizing the bystander MN1 axon in different genotypes: control, phasic MN-Is ablated (*Is>RPR*), tonic MN2, MN9, MN10 ablated (*CQ>RPR*), and both tonic and phasic MNs ablated (*Is ∪ CQ>RPR*). Schematics (left) depict ablated MNs and their effects on MN1. In controls, MN1 exclusively innervates muscle 1 (top row), and this pattern is unchanged after phasic MN-Is ablation (second row). Ablation of tonic MNs (CQ>RPR) induces sprouting of MN1 onto muscle 2 (M2). Co-ablation of tonic and phasic MNs (*Is ∪ CQ>RPR*) induces robust MN1 sprouting, forming ectopic NMJs on all nearby denervated muscles (M2, M9, M10) in addition to its native target (M1). Postsynaptic sites are marked by Dlg (magenta) and muscles by phalloidin (blue). MN1 is labeled with UAS-myr-GFP driven by 94G06- Gal4; targeted MN ablation is achieved using the LexA/OP-RPR system. (E) Quantification of MN1 axon and NMJ parameters across groups shown in (D). (i) Total MN1 area on DL muscles 1, 2, 9, and 10. *Is ∪ CQ>RPR* larvae exhibit significantly increased total MN1 surface area relative to control and other ablation groups. (ii) Total number of MN1 boutons on DL muscles. No significant differences across groups. (iii) Average bouton size (area/bouton number). No significant differences across groups. (iv) Proportional distribution of MN1 NMJ area on individual DL muscles. In CQ>RPR larvae, MN1 ectopically innervates muscle 2; in *Is ∪ CQ>RPR* larvae, MN1 innervates muscles 2, 9, and 10 in addition to its native target Muscle 1. The area on Muscle 1 is significantly decreased compared to control group. (v) Average number of motor axon branching points (green) and NMJ branching points. *CQ>RPR* and *Is ∪ CQ>RPR* groups exhibit significantly increased axon branching compared to control group. For control, *Is>RPR*, and *Is ∪ CQ>RPR*, n = 8 larvae (16 hemisegments). For *CQ>RPR*, n = 7 larvae (13 hemisegments). Data points represent single hemisegments. Error bars: ±SEM. Statistical analysis: Kruskal-Wallis with Dunn’s post-hoc test (*p<0.05; **p<0.01; ***p<0.001; \*\*\**p<0.0001*).

In this study, we focused on the dorsal-longitudinal (DL) muscles, specifically Muscles 1, 2, 9, and 10 (**Fig. 1A**). These muscles receive inputs from a common phasic-firing MN- Is, also known as RP2 or the common dorsal excitor (CDE), and four distinct tonic-firing (Ib) MNs—MN1, MN2, MN9, and MN10—each innervating an individual muscle (**Fig. 1A**) (5, 30–36).

Previous studies have shown that apoptotic ablation of the phasic MN-Is (hereafter referred to as **Is**) leads to structural expansion of the neuromuscular junction (NMJ) formed by the co-innervating tonic MN1, increasing tonic input to Muscle 1. In contrast, ablation of the tonic MN1 does not trigger structural changes in the NMJ formed by MN- Is (7–10). These findings highlight intrinsic differences in how tonic and phasic MNs respond to the loss of their co-innervating partners. However, these studies removed only one MN class—either tonic or phasic—leaving the target muscles partially innervated.

Thus, it remained unclear whether complete removal of both excitatory inputs could trigger compensatory sprouting by other MNs. To address this, we genetically ablated either or both tonic and phasic MNs innervating Muscles 2, 9, and 10 and examined structural remodeling by the bystander MN1 (**Fig. 1B–D**). For ablation, we used previously validated MN-LexA drivers (37, 38) to express the pro-apoptotic genes *reaper (rpr)* and *head involution defective (hid)*, inducing cell death in targeted MNs (39). The tonic MN1 was visualized using 94G06 (40) (MN1 Ib)- Gal4 driving UAS-myr-GFP.

We found that MN1 did not form ectopic NMJs when only the phasic MN-Is was ablated (Is >RPR) and only formed ectopic NMJs on Muscle 2 when tonic MNs 2, 9, and 10 were ablated (CQ >RPR) (**Fig. 1 D,E**). In contrast, concurrent ablation of both tonic and phasic MNs (CQ ∪ Is > RPR) triggered robust sprouting of MN1, which extended axons and formed de novo NMJs on fully denervated Muscles 2, 9, and 10 (**Fig. 1 D,E**).

Quantitative morphometric analysis (41, 42) (**Fig. 1 C)** revealed that MN1 in CQ ∪ Is >RPR larvae showed a significant increase in overall NMJ area across all four DL muscles (1, 2, 9, and 10) compared to controls, while average bouton size and total bouton number remained unchanged (**Fig. 1 D,E**). Importantly, the total MN1 NMJ area was redistributed across the DL muscles, with reduced allocation to its native target Muscle 1 compared to controls—suggesting an upper limit on the total synaptic area MN1 can establish (**Fig. 1 D,E**). Finally, we quantified total axonal and NMJ branch points. CQ > RPR and CQ ∪ Is > RPR larvae showed a significant increase in axon branch points (GFP-only), consistent with a sprouting phenotype, whereas NMJ branch points (Dlg+GFP) were not significantly altered (**Fig. 1 D,E**).

Together, these results demonstrate that robust heterosynaptic sprouting of MN1 is selectively triggered by complete loss of both tonic and phasic excitatory inputs to neighboring muscles.

### Phasic MN-Is Maintains Its Normal Innervation After Ablation of Co-innervating Tonic MNs

Our earlier experiments demonstrated that the tonic MN1 exhibits robust heterosynaptic sprouting when neighboring muscles are fully deprived of both tonic and phasic inputs. We next asked whether the phasic MN-Is possesses a comparable capacity for structural remodeling and NMJ expansion following the loss of its tonic co-innervators.

Previous studies, in which only MN1 was ablated, reported no structural changes in NMJs formed by the phasic MN-Is on Muscle 1 (7–10). To extend these findings, we ablated additional tonic MNs (MN1, MN2, MN9, and MN10), thereby fully depriving the dorsal-longitudinal (DL) muscles 1, 2, 9, and 10 of all tonic innervations (**Fig. 2A**). In both control and ablated groups, the phasic MN-Is was visualized using myr-GFP driven by an enhancer element cloned from the 27E09-Gal4 driver, which selectively targets phasic MNs.

**Figure 2.**
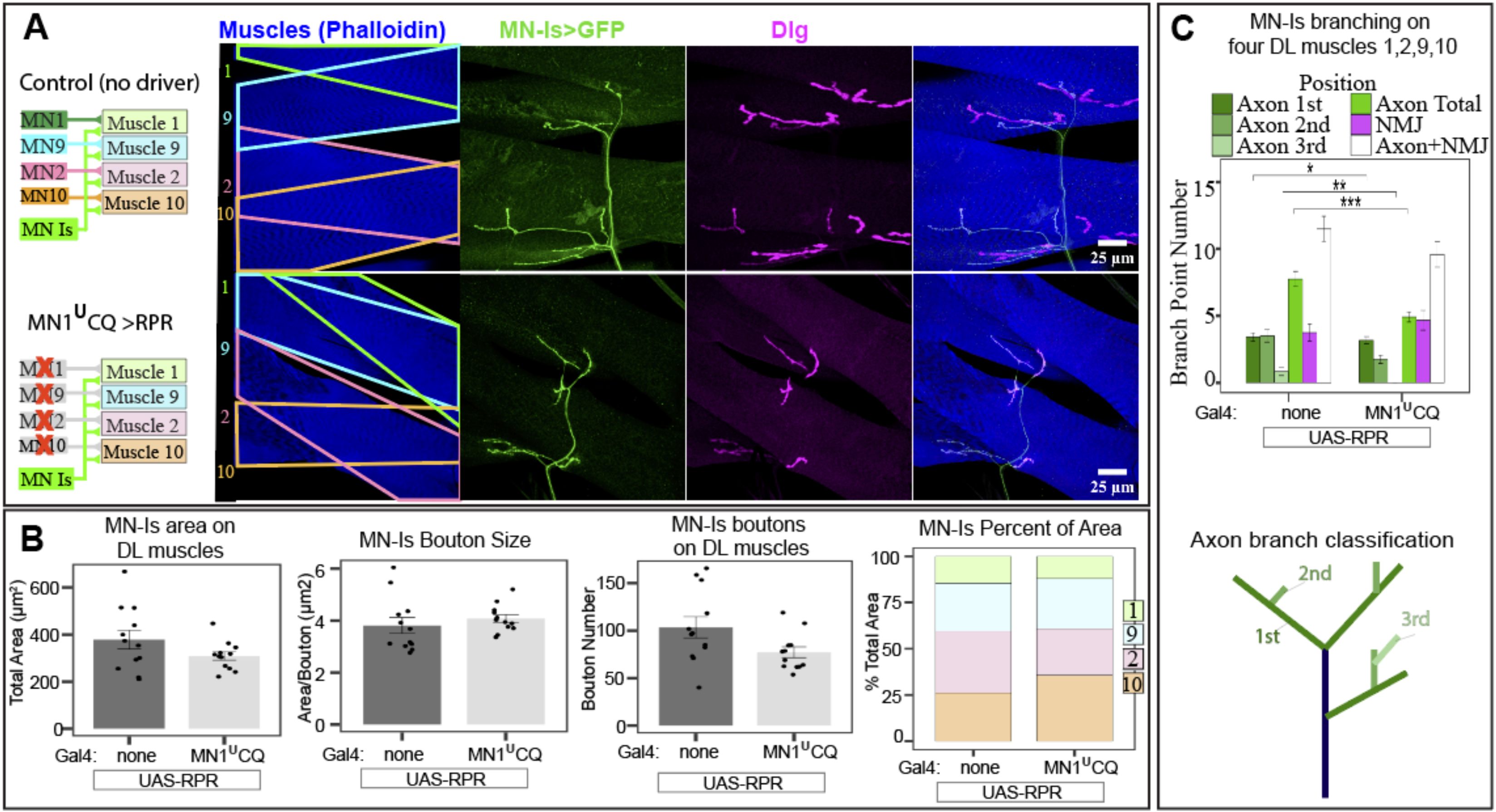
Phasic MN-Is does not exhibit structural expansion following ablation of tonic MNs innervating dorsal-longitudinal (DL) muscles. (A) Representative NMJ immunostaining showing the bystander phasic MN-Is axon in control larvae and in larvae where all tonic Ib MNs innervating DL muscles (MN1, MN2, MN9, MN10) were ablated (*MN1 ∪ CQ>RPR*). Postsynaptic sites are labeled with Dlg (magenta), muscles with phalloidin (blue), and MN-Is axons with myr-GFP driven by the 27E09 enhancer. Ablation of tonic MNs was achieved using the 94G06-Gal4 ∪ CQ-Gal4 / UAS-RPR system. (B) Quantification of Is NMJ parameters shows no significant differences in total NMJ area, bouton size, bouton number, or NMJ area distribution between control and *MN1 ∪ CQ>RPR* larvae. Each data point represents a single hemisegment. (C) Ablation of tonic MNs results in a significant reduction in 2nd- and 3rd-order axonal branches of the bystander MN-Is. For both control and *MN1 ∪ CQ>RPR* groups, n = 6 larvae (12 hemisegments). Data points represent individual hemisegments. Error bars indicate mean ± SEM. Statistical analysis: Kruskal-Wallis with Dunn’s post-hoc test (*p*<0.05; \**p*<0.01; \*\**p*<0.001; \*\*\**p*<0.0001).

Following tonic MN ablation (CQ ∪ MN1 >RPR), the intact phasic MN-Is retained its normal innervation pattern on Muscles 1, 2, 9, and 10 (**Fig. 2A**). Apart from a modest reduction in the number of 2nd- and 3rd-order axonal branches in CQ ∪ MN1 >RPR larvae, there were no significant differences in bouton number, area, or size between control and ablated groups (**Fig. 2B, C**). Despite complete loss of tonic excitatory input, the phasic MN-Is did not exhibit ectopic sprouting or NMJ expansion on these muscles.

This lack of response contrasts sharply with the extensive sprouting observed in MN1 under similar conditions, underscoring a fundamental difference in the plasticity potential of tonic versus phasic MNs. Consistent with earlier studies (7–10), these results suggest that phasic MN-Is lacks the intrinsic capacity for compensatory structural remodeling following the loss of its co-innervating tonic partners.

### MN1 Sprouting is Induced by Ablation but Not by Chronic or Acute Silencing of Neighboring MNs

To confirm that the robust sprouting observed in MN1 results specifically from the loss of neighboring motor neurons—and not from artifacts of the LexA/AOP genetic system—we repeated the ablation experiments using the Gal80^ts^/GAL4/UAS system. Gal80^ts^ inhibits GAL4 activity at 24°C but not at 29°C (43, 44).

Using previously characterized CQ and Is GAL4 drivers (7, 36–38, 40, 45), we targeted the tonic MN2, MN9, and MN10 along with the phasic MN-Is for UAS-reaper (rpr)- mediated ablation. MN1 was labeled with MN1-lexA >AOP-myr-GFP. Animals raised continuously at 29°C showed robust MN1 sprouting and ectopic NMJs on denervated Muscles 2, 9, and 10, whereas controls maintained at 24°C (with functional Gal80 protein blocking GAL4 activity) showed no sprouting (**Fig. 3A-B, Supp Fig S1**). These findings confirm that MN1 sprouting is specifically triggered by the physical loss of neighboring MNs rather than by potential genetic artifacts.

**Figure 3.**
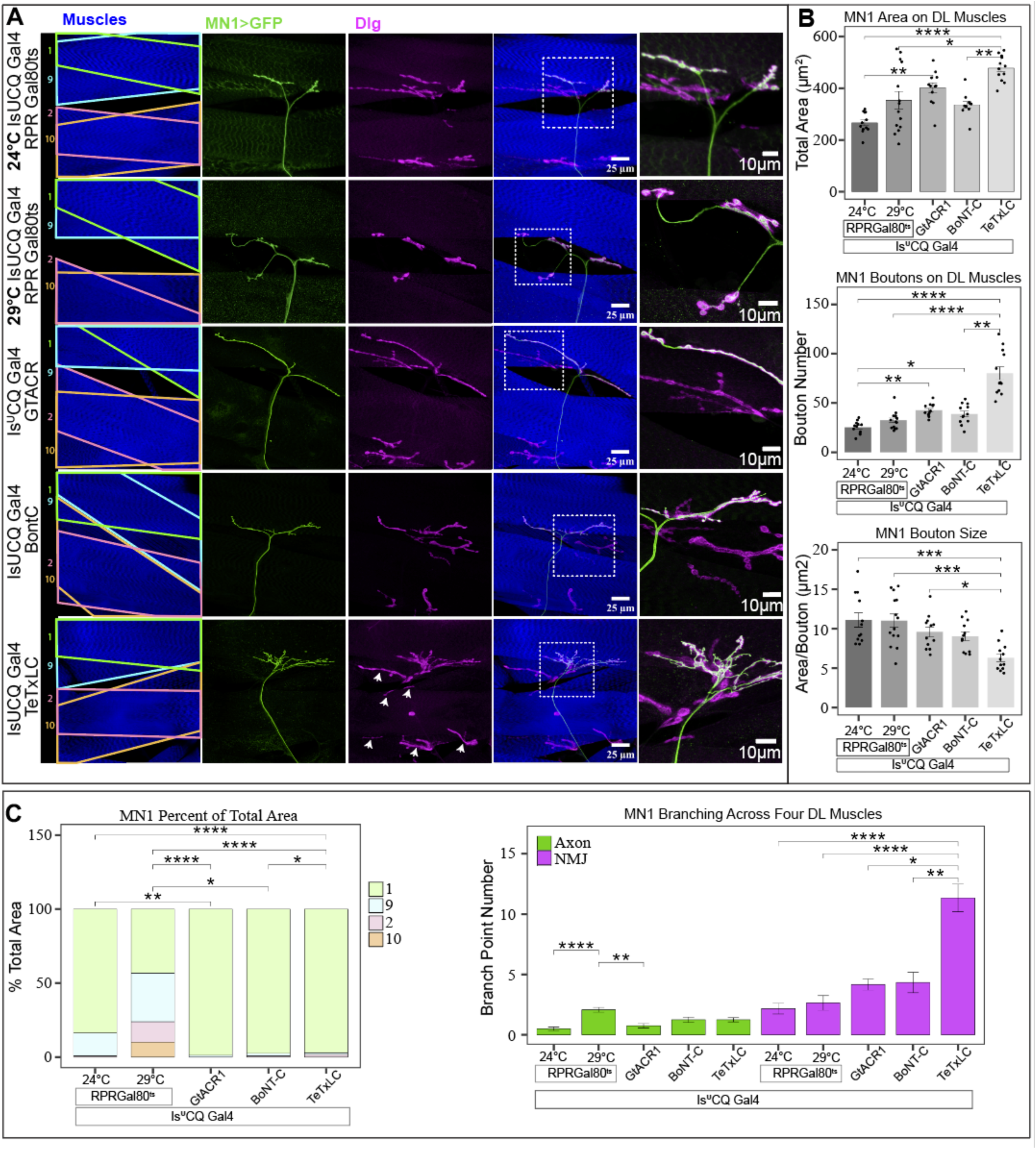
The bystander MN1 exhibits distinct structural responses to different perturbations of neighboring MNs. (A–C) Representative images (A) and quantitative analyses (B–C) of MN1 axons and NMJs across genotypes. MN1 is labeled with MN1-lexA >AOP-myr-GFP. In control animals (CQ ∪ Is-Gal4/UAS-RPR, Gal80^ts^ raised at 24°C), MN1 shows normal axon and NMJ morphology. In the ablation group (CQ ∪ Is-Gal4/UAS-RPR, Gal80^ts^ raised at 29°C), MN1 exhibits robust sprouting, forming ectopic NMJs on denervated muscles 2, 9, and 10. In the acute silencing group (CQ ∪ Is-Gal4/UAS-GtACR1), MN1 NMJ area and bouton number are elevated relative to controls, but no ectopic sprouting onto muscles 2, 9, and 10 is observed. In the chronic silencing group targeting both evoked and spontaneous release (CQ ∪ Is- Gal4/UAS-BoNT-C), MN1 shows increased bouton numbers without ectopic sprouting. In the chronic silencing group targeting only evoked release (CQ ∪ Is-Gal4/UAS-TeTxLC), MN1 NMJ branching, area, and bouton number are markedly increased compared to controls and other perturbations. However, bouton size is significantly reduced, and no ectopic sprouting onto muscles 2, 9, and 10 occurs. Notably, the overall NMJ morphology of TeTxLC-silenced tonic and phasic MNs (white arrows) appears normal based on Dlg staining. Data points represent single hemisegments. Ablation group: n=7 larvae (14 hemisegments). All other groups: n=6 larvae (12 hemisegments). Error bars: ±SEM. Kruskal-Wallis with Dunn’s post-hoc test (*p<0.05; **p<0.01; ***p<0.001; ****p<0.0001).

We next tested whether silencing the activity of neighboring MNs—without their physical removal—would be sufficient to induce MN1 sprouting. For acute silencing, we used the light-activated chloride channel GtACR1, which causes chloride influx and neuronal hyperpolarization upon light exposure (46, 47). Previous works have demonstrated the efficacy of GtACR1 in silencing MNs and other neurons in Drosophila larvae (37, 45, 48, 49). In CQ ∪ Is >GtACR1 larvae, MN1 maintained its normal innervation of Muscle 1 and did not form ectopic NMJs on Muscles 2, 9, or 10. However, we observed a modest increase in MN1 NMJ area and bouton number compared to controls, without changes in bouton size (**Fig. 3A-B, Supp Fig S1**).

We then tested chronic silencing using botulinum neurotoxin-C (BoNT-C), which blocks both evoked and spontaneous neurotransmitter release (27, 28). In CQ ∪ Is >BoNT-C larvae, MN1 did not form ectopic NMJs, but the number of boutons on Muscle 1 was significantly increased relative to controls, despite no changes in NMJ area or bouton size. (**Fig. 3A-B Supp Fig S1**).

We then expressed tetanus toxin light chain (TeTxLC), which blocks evoked but not spontaneous neurotransmitter release (50). In CQ ∪ Is >TeTxLC larvae, MN1 again did not form ectopic NMJs, but showed substantial NMJ expansion on its native target, Muscle 1 (**Fig 3 A-C Supp Fig S1**). This included significantly increased NMJ area, branching, and bouton number, along with a marked reduction in average bouton size (**Fig 3 B-C Supp Fig S1**). These features are consistent with satellite bouton formation previously observed following TeTxLC-mediated silencing of MN-Is (7, 8). Notably, Dlg staining revealed that the NMJs of TeTxLC-silenced tonic MNs maintained normal morphology and did not exhibit the extensive NMJ overgrowth observed in the bystander MN1 (**Fig. 3A)**, suggesting that the sprouting phenotype is specific to the unaffected neuron responding to loss of presynaptic activity in neighboring inputs.

Taken together, these results demonstrate that heterosynaptic sprouting of MN1 requires the physical ablation of neighboring MNs. In contrast, silencing of neighboring MNs— whether acute or chronic—does not trigger ectopic sprouting, though it can modulate bouton number and NMJ structure at the native target.

### Ectopic MN1 Sprouting Restores Muscle Activation and Partially Rescues Crawling Behavior

Thus far, we have shown that concurrent ablation—but not silencing—of neighboring tonic and phasic MNs triggers robust sprouting of the bystander MN1, resulting in ectopic NMJs on fully denervated Muscles 2, 9, and 10. To determine whether these ectopic NMJs are functional and capable of activating the muscles they innervate, we performed in vivo muscle calcium imaging in intact, forward-crawling larvae.

As a baseline, we first examined muscle calcium activity in larvae where all tonic and phasic MNs innervating dorsal muscles 1, 2, 9, and 10 were either genetically ablated (CQ-lexA, 27E09-lexA; 94G06-lexA >RPR) or acutely silenced (CQ-lexA, 27E09-lexA; 94G06-lexA >GtACR1) (**Fig 4 A-C**). In both groups, the dorsal-longitudinal (DL) muscles failed to show calcium transients or contraction during forward crawling, confirming complete loss of excitatory input. In contrast, ventral and lateral muscles with intact MN innervation exhibited robust calcium activity and contractions (**Fig 4 A,B; Video S1, Video S2, Video S3**). These observations validate that (i) both GtACR1- mediated silencing and RPR-mediated ablation effectively eliminate MN output and (ii) the MN driver lines selectively target the intended MNs without affecting off-target MNs.

**Figure 4.**
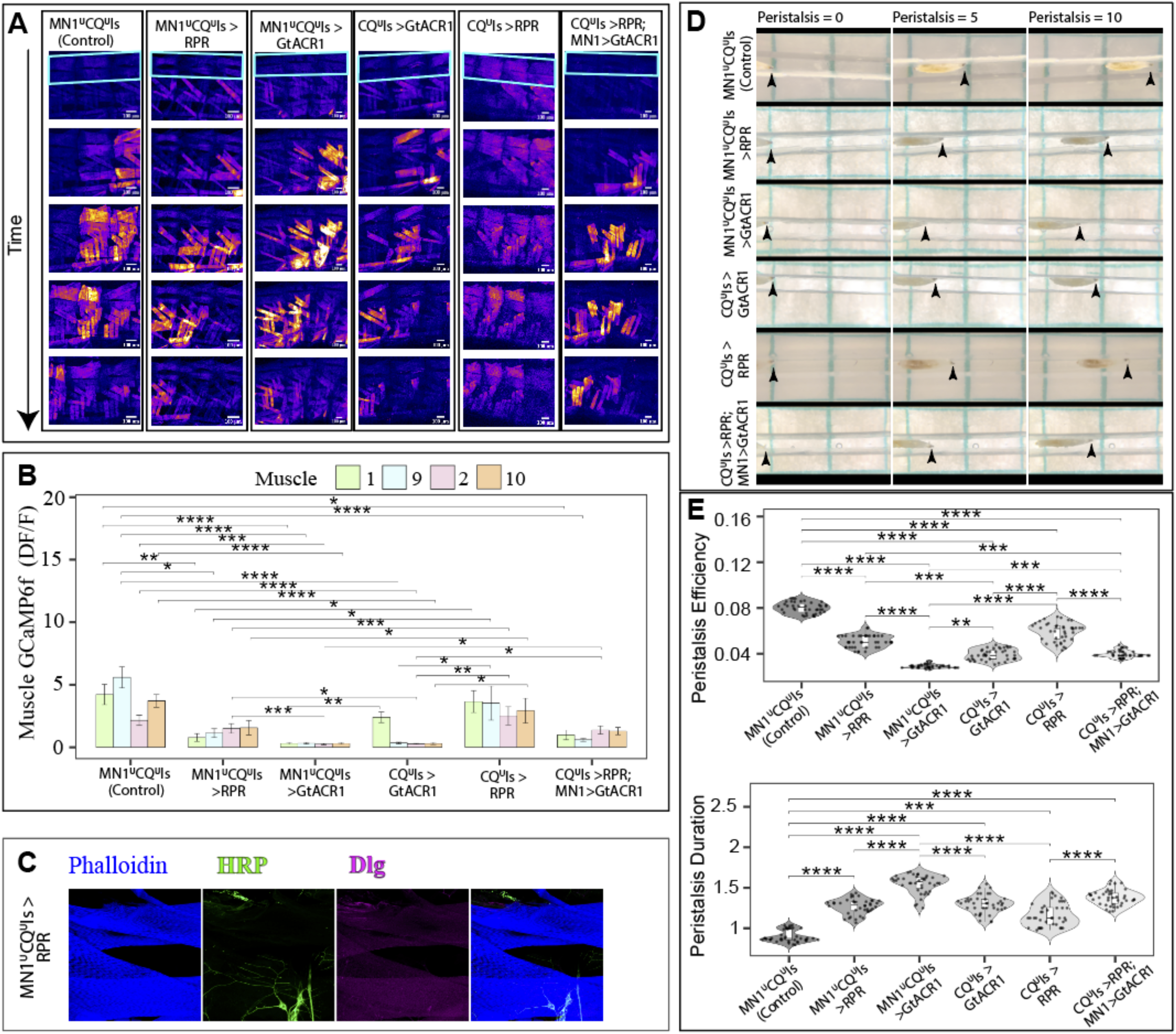
Ectopic NMJs formed by sprouting MN1 are functional and improve larval locomotion. (A) Muscle calcium imaging in intact, forward-crawling larvae of different genotypes. Each column shows still images from three adjacent abdominal segments at successive time points during a forward peristaltic wave. GCaMP6f fluorescence (expressed under the 44H10 muscle-specific enhancer) increases during muscle contraction. Cyan rectangles mark dorsal-longitudinal (DL) muscles in the top panels. In control larvae (MN1 ∪ CQ ∪ Is-lexA/ Aop-GTACR ATR (-)), DL muscles contract sequentially, with GCaMP6f signals propagating from posterior (right) to anterior (left) during crawling. Ablation of all tonic and phasic MNs (MN1 ∪ CQ ∪ Is-lexA/Aop-RPR) abolishes DL muscle activity, leaving ventral and lateral muscles unaffected. Similarly, acute silencing of these MNs (MN1 ∪ CQ ∪ Is-lexA/Aop-GtACR1 ATR (+)) eliminates DL muscle activity. In CQ ∪ Is-lexA/Aop-GtACR1 ATR (+) larvae, only Muscle 1 remains active, consistent with preserved MN1 input. In contrast, CQ ∪ Is-lexA/Aop-RPR larvae exhibit robust calcium activity and contractions in all four DL muscles, indicating functional reinnervation by sprouted MN1. Acute silencing of the sprouted MN1 in CQ ∪ Is- lexA/Aop-RPR; MN1-Gal4>UAS-GtACR1 ATR (+) larvae abolishes DL muscle activity. Frames are taken from Supplementary Video X. (B) Quantification of GCaMP6f signal in DL muscles (1, 2, 9, and 10) confirms observations in (A). n = 5 larvae per genotype (10 hemisegments total). Statistical analysis: Kruskal–Wallis test with Dunn’s post-hoc correction (*p<0.05, **p<0.01, ***p<0.001, ****p<0.0001). (C) Immunostaining of DL muscles in MN1 ∪ CQ ∪ Is-lexA/Aop-RPR larvae verifies complete ablation of targeted MNs. Anti-HRP (green) labels the nearby sensory neurons, and no motor axons or Dlg puncta are detected in denervated DL muscles. (D) Representative crawling trajectories of larvae from the genotypes in (A). Each row shows larval positions at the start line (Peristalsis = 0), after 5 peristalses, and after 10 peristalses. Among all groups, control larvae (MN1 ∪ CQ ∪ Is-lexA/+) ranked first in distance traveled, followed by CQ ∪ Is-lexA/Aop-RPR larvae, which ranked second. In contrast, MN1 ∪ CQ ∪ Is-lexA/Aop-GtACR1 larvae showed the least forward progression. (E) Quantification of peristalsis efficiency (distance traveled per peristaltic wave) and peristalsis duration (time required to complete a wave). Control larvae (MN1 ∪ CQ ∪ Is- lexA/+) exhibited the highest peristalsis efficiency and shortest duration, followed by CQ ∪ Is-lexA/Aop-RPR larvae. Silencing the sprouted MN1 in CQ ∪ Is-lexA/Aop-RPR; MN1-Gal4>UAS-GtACR1 larvae reduced locomotor performance to levels comparable with other two silenced groups. n = 40 larvae per group, with each dot representing an individual animal. Statistical analysis: One-way ANOVA with Tukey post-hoc test (*p<0.05, **p<0.01, ***p<0.001, ****p<0.0001).

We then repeated these experiments but spared MN1. In the acute silencing group (CQ-lexA, 27E09-lexA >GtACR1), only Muscle 1 displayed calcium transients and contraction during crawling, consistent with MN1 maintaining its normal innervation (**Fig 4 A,B; Video S4**). By contrast, in the ablation group (CQ-lexA, 27E09-lexA >RPR), all four dorsal muscles (1, 2, 9, and 10) exhibited calcium transients and contraction, demonstrating that ectopic NMJs formed by MN1 are capable of restoring contractile activity in denervated muscles (**Fig 4 A,B; Video S5**).

To directly test whether the sprouted MN1 drives these responses, we acutely silenced MN1 (94G06-Gal4 >GtACR1) in larvae where neighboring MNs had been ablated (CQ-lexA, 27E09-lexA >RPR). Silencing MN1 abolished calcium activity in all four dorsal muscles during crawling, confirming that the sprouted MN1 terminals are functionally active and required for muscle contraction (**Fig 4 A,B; Video S6**).

Next, we investigated whether MN1 sprouting contributes to behavioral recovery. We quantified forward crawling by measuring peristalsis efficiency (distance traveled per peristalsis) and peristalsis duration (time to complete a forward wave). Control larvae displayed a peristalsis efficiency of ∼0.8 mm per wave. Larvae lacking dorsal MN inputs (CQ-lexA, 27E09-lexA; 94G06-lexA >RPR) showed markedly reduced efficiency, but those with intact MN1 (CQ-lexA, 27E09-lexA >RPR) performed significantly better, indicating partial restoration of crawling (**Fig 4 D,E; Video S7**).

Notably, CQ-lexA, 27E09-lexA >RPR larvae also outperformed CQ-lexA, 27E09-lexA >GtACR1 larvae (**Fig 4 D,E; Video S7**), suggesting that ablation-induced sprouting of MN1 (occurring in the ablated group but not in the silenced group) underlies this improvement. However, because 27E09-lexA also targets ventral-projecting phasic MNs, it is possible that structural plasticity of ventral tonic MNs contributed to recovery. To isolate MN1’s contribution, we silenced the sprouted MN1 in CQ-lexA, 27E09-lexA >RPR larvae using 94G06-Gal4 >GtACR1. This manipulation significantly reduced peristalsis efficiency compared to unsilenced CQ-lexA, 27E09-lexA >RPR larvae (**Fig 4 D,E; Video S7**), directly demonstrating that MN1 sprouting is adaptive and improves locomotion.

Across all genotypes, we observed an inverse relationship between peristalsis efficiency and peristalsis duration: larvae traveling shorter distances per peristaltic wave also exhibited slower propagation of contractions across body segments, resulting in longer peristalsis durations (**Fig 4 E; Video S7**).

Together, these results establish that MN1 ectopic sprouting restores muscle contractility and partially rescues motor behavior, highlighting its role as a compensatory response following injury.

### Ectopic MN1 Sprouting Occurs Primarily During Larval Stages

In *Drosophila*, embryogenesis lasts ∼22–24 hours at 25°C, after which the embryo hatches into a first instar larva (L1) capable of foraging, feeding, and performing locomotor behaviors such as forward crawling. By hatching, core motor circuits are already established, with motor neurons forming functional synapses onto their target muscles. The L1 stage lasts ∼25 hours, followed by molting into second (L2) and third instar (L3) stages, each lasting ∼24–25 hours before pupation. Our initial analyses of MN1 sprouting and ectopic NMJ formation were performed in L3 larvae. To better define the timeline of these structural changes, we examined MN1 remodeling across the L1–L3 transition in animals with ablated tonic and phasic MNs.

Using live imaging of MN1 labeled with tdTomato, we tracked axonal branching in individual larvae at L1, L2, and L3 stages. Between imaging sessions, larvae were recovered and returned to food to continue development. This allowed us to quantify and distinguish 1st-, 2nd-, and 3rd-order axon branches extending from MN1’s main shaft (**Fig. 5A schematic**).

**Figure 5.**
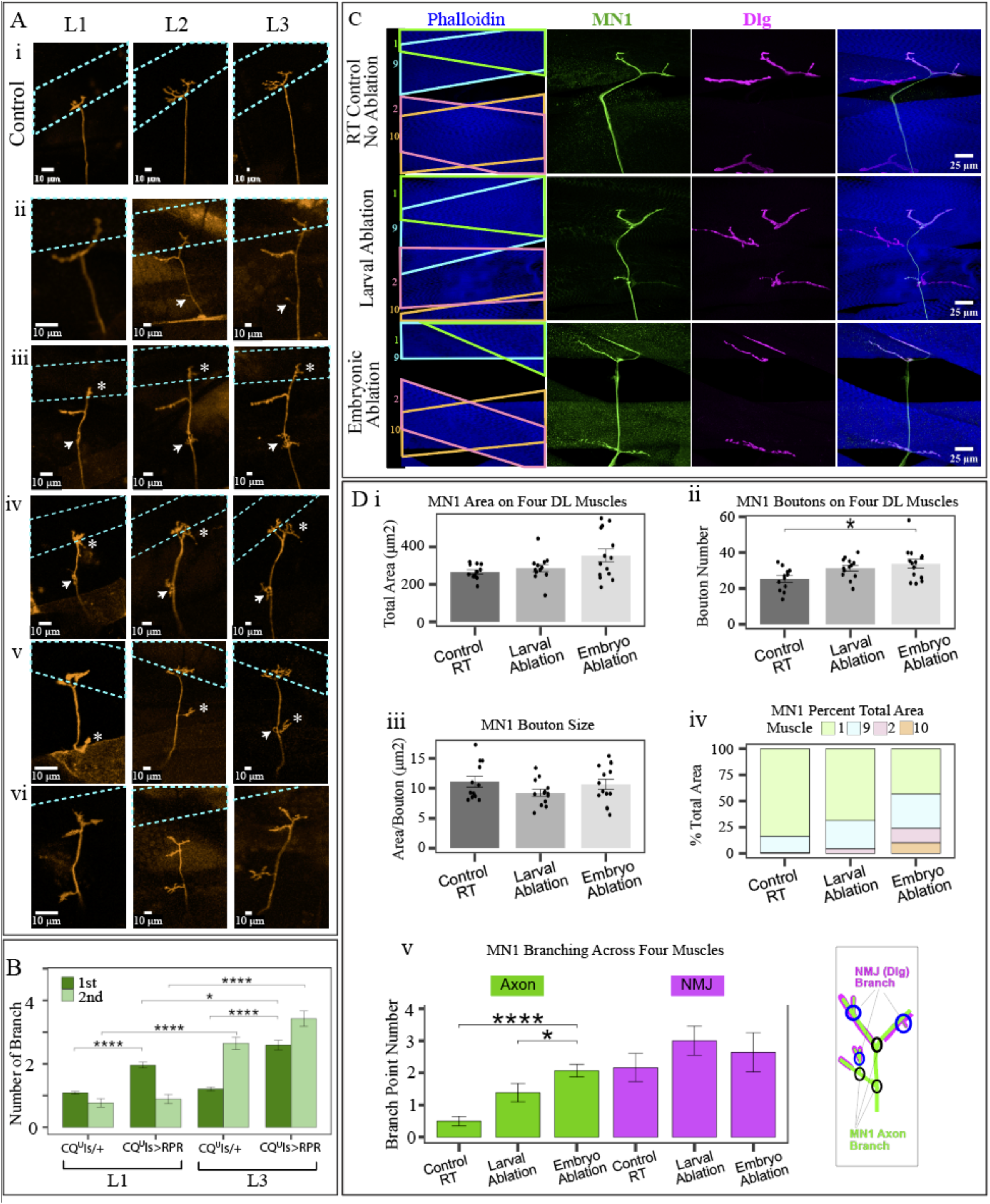
MN1 axon sprouting dynamics and timing of injury-induced plasticity (A) Live imaging of MN1 axons across larval stages. Each row shows stills from the same hemisegment at L1, L2, and L3. (i) In a control larva (CQ ∪ Is-Gal4/+; MN1-lexA >Aop-tdTomato), MN1 innervates its native target muscle 1 (cyan box) by L1. The axon terminal arborizes further between L1–L3, with the main shaft bifurcating beneath muscle 1 to form a T-shaped NMJ. (ii–vi) In ablated larvae (CQ ∪ Is-Gal4/UAS-RPR; MN1-lexA >Aop-tdTomato), diverse sprouting patterns are observed: (ii) New axon branches (white arrow) emerge at L2 and grow further by L3. (iii–v) Small axon buds appear at L1 (white arrows) and expand progressively in L2 and L3. Some branches (white asterisks) are already present at L1 and undergo marked growth during later stages. (vi) In rare cases, MN1 fails to reach its native target muscle 1 and instead branches prematurely to innervate other DL muscles. (B) Quantification of 1st- and 2nd-order axon branches in L1 and L3 stages for controls and ablated animals. At L1, ablated larvae show ∼2× more 1st-order branches than controls, with comparable 2nd-order branching. By L3, ablated larvae exhibit significantly more 1st-order branches than controls. Within ablated larvae, both branch types increase markedly from L1 to L3; in controls, only 2nd-order branching increases over time. Data points represent single hemisegments (L1 control: n=5 larvae, 57 hemisegments; L1 ablation: n=5 larvae, 56 hemisegments; L3 control: n=5 larvae, 57 hemisegments; L3 ablation: n=5 larvae, 54 hemisegments). Error bars: ±SEM. Kruskal-Wallis with Dunn’s post-hoc test (*p<0.05; **p<0.01; ***p<0.001; ****p<0.0001). (C) Timing of ablation influences sprouting. Post-embryonic (larval) ablation of MNs (CQ ∪ Is-Gal4/UAS-RPR,Gal80^ts^ shifted from 24°C to 29°C at L1) still triggers MN1 sprouting but less robustly than embryonic ablation (CQ ∪ Is-Gal4/UAS-RPR,Gal80^ts^ raised at 29°C throughout). No sprouting occurs in control larvae maintained at 24°C. (D) Quantification of MN1 bouton size, bouton number, axon area, and branch complexity across groups in (C). Data points represent single hemisegments. Control group: n = 6 larvae (12 hemisegments). For embryonic ablation: n = 7 larvae (14 hemisegments). Embryonic and control datasets are the same as in Fig. 3. For larval ablation, n = 7 larvae (13 hemisegments). Error bars: ±SEM. Kruskal-Wallis with Dunn’s post-hoc test (*p<0.05; **p<0.01; ***p<0.001; ****p<0.0001). encing lsuca Gal4>BontC

At L1, MN1 axons in ablated larvae already exhibited a twofold increase in 1st-order branches compared to controls, while 2nd-order branching was comparable between groups. By L3, ablated larvae showed significantly higher numbers of 1st-order branches relative to controls. Within the ablated group, both 1st- and 2nd-order branch numbers increased significantly from L1 to L3, whereas in controls only 2nd-order branching increased across development. These findings suggest that while MN1 sprouting in ablated animals is initiated early, the majority of axon branching and ectopic NMJ formation occurs progressively during the L1–L3 transition **(Fig. 5A–B; Supp. Fig.)**.

To test whether MN1 sprouting can be induced exclusively during larval stages, we performed temporally restricted ablation. Since MN-Gal4 drivers (e.g., CQ-Gal4, 27E09- Gal4) activate UAS-rpr/hid from embryogenesis onward, we used Gal80^ts^ to suppress Gal4 activity at 24°C and permit it at 29°C (43, 44). For larval-only ablation, embryos were maintained at 24°C until hatching and then shifted to 29°C from L1 through L3 prior to NMJ analysis. As reference points, we used data from embryonic ablation (continuous 29°C) and control animals (continuous 24°C) described earlier in **Fig. 3**. Larvae subjected to larval-stage ablation showed MN1 sprouting and ectopic NMJ formation, though to a lesser extent than embryonically ablated animals (**Fig. 5C–D**).

Taken together, these results indicate that MN1 sprouting is initiated early but progresses substantially throughout larval development, with the L1–L3 period representing a key window for ectopic NMJ formation following injury.

## Discussion

### Subtype-Specific Plasticity Revealed Through Complete Denervation

Our study reveals a robust, subtype-specific capacity for structural plasticity within an established motor circuit following injury. We demonstrate that tonic MN1, but not phasic MN-Is, undergoes extensive axonal sprouting and ectopic NMJ formation when neighboring tonic and phasic MNs are ablated. These ectopic NMJs restore muscle contractility and contribute to improved locomotor performance, providing functional evidence that the observed plasticity is adaptive. In contrast, when neighboring MNs are functionally silenced without ablation, MN1 does not sprout, indicating that complete physical loss—not merely loss of synaptic activity—is required to trigger heterosynaptic structural plasticity (HTSP).

Our approach builds on earlier observations by Chang and Keshishian (1996), who reported that embryonic denervation could lead to some ectopic innervation from nearby neurons (51). However, they utilized laser-mediated MN ablation, which resulted in variable levels of denervation and often lacked control over which neurons contributed to reinnervation. In contrast, we used targeted genetic ablation to achieve reproducible, complete denervation of specific muscles, revealing that sprouting arises specifically from the identified tonic MN1. Furthermore, we show that the newly formed NMJs are functional and that sprouting mainly occurs after larval hatching and continues across larval stages, defining a broader temporal window for plasticity.

In more recent studies investigating NMJ plasticity, either tonic or phasic MNs were disrupted, but never both. As a result, target muscles continued to receive partial excitatory input from surviving MNs, allowing for homeostatic compensatory changes within an already innervated context. Thus, the effects of completely removing all excitatory input from a given muscle and testing whether other MNs can sprout to reinnervate these fully denervated targets remained unknown. To address this gap, we developed genetic tools enabling the simultaneous ablation of both tonic and phasic MNs innervating specific muscles, creating a fully denervated environment. This strategy revealed a previously unrecognized capacity for bystander MN1 to form de novo ectopic NMJs and reestablish motor output. This subtype specificity aligns with prior findings that tonic Ib MNs—but not phasic Is MNs—exhibit structural and functional compensation following input loss (7, 8, 10). Our findings extend this framework by demonstrating that MN1 can not only expand existing synaptic terminals but also innervate novel postsynaptic targets, forming functional ectopic NMJs on muscles it does not typically contact. This level of remodeling goes beyond classic homeostatic plasticity and reflects a more profound form of structural reorganization.

Importantly, MN1 sprouting occurred only following the physical ablation of neighboring MNs and not when those MNs were chronically or acutely silenced. Our findings further support that tonic MNs possess a greater intrinsic capacity for structural remodeling in response to injury compared to phasic MNs. This finding agrees with Han et al. (2022), who showed that botulinum toxin silencing—even when eliminating both evoked and spontaneous transmission—does not induce structural compensation, whereas physical loss does (27). These results emphasize that circuit rewiring is not merely a response to inactivity but likely requires injury-associated cues such as axonal degeneration or glial signaling.

Our injury model appears to diverge from CNS critical period constraints. Ackerman et al. (2021) silenced MN1 and MN-Is and examined dendritic remodeling within the same MNs, identifying an early embryonic window during which activity loss induces dendritic growth (52). In contrast, we demonstrate that MN1 can undergo peripheral axonal remodeling beginning 0–4 h after larval hatching and continuing throughout larval stages, highlighting that different compartments of the same neuron (dendrites vs. axons) follow distinct temporal rules for plasticity.

### Intrinsic Differences in Plasticity Between Tonic MN1 and Phasic MNs

MN1 displays a markedly greater intrinsic capacity for structural plasticity compared to its phasic counterparts, such as the phasic MN-Is. This enhanced plasticity is likely underpinned by a multilayered set of molecular, structural, genetic, and functional differences at the synaptic, transcriptional, and cellular level (28, 38, 53, 54).

Structurally, tonic MN1 forms larger synaptic boutons with expansive, donut-shaped active zones and a 4.4-fold larger neuromuscular junction area at muscle 1 containing 3.8-fold more active zones (AZ), creating an ideal substrate for growth and remodeling, unlike the compact active zones of phasic neurons optimized for rapid transmission rather than plasticity (13, 27, 28, 38, 54). Functionally, tonic MN1 displays lower basal synaptic output and more "silent" active zones possibly conducive to sprouting, whereas phasic MN-Is show high initial release probability (Pr) that depresses with repetitive stimulation (54). Importantly, high-Pr phasic MN-Is synapses show denser voltage-gated calcium channel (VGCC) clustering and tighter **vesicle** coupling—architectural features that could enhance release efficiency but may potentially constrain structural remodeling and growth (54).

At the molecular level, MN1 expresses significantly elevated levels of growth-promoting genes including Wnt4 (40-fold higher), Pyramus, Toll-6, and SiaT—establishing a growth-permissive transcriptional program and environment (38). Conversely, phasic neurons upregulate calcium buffering proteins like Cbp53E (30-fold higher) and vesicle trafficking chaperones that constrain remodeling (38). MN1’s superior calcium handling—maintaining higher resting calcium levels while enabling rapid extrusion after activity—prevents growth inhibition and supports sprouting, unlike phasic neurons that accumulate excessive calcium during stimulation with slower recovery. Furthermore, MN1’s distinct post-translational modifications, cytoskeletal regulation, and broader distribution of scaffolding proteins like Bruchpilot could likely create a cellular environment permissive to structural reorganization and functional recovery after injury, fundamentally distinguishing it from phasic neurons that are intrinsically optimized for high-fidelity transmission rather than adaptive structural change.

### Evolutionary Parallels in Injury-Induced Plasticity

Our findings reflect a conserved strategy for injury-induced plasticity across species. In the mammalian spinal cord, both interneurons and descending corticospinal tract (CST) neurons can sprout following injury, particularly when targets are fully or partially denervated (55). Similar to our findings in Drosophila, this rewiring often originates from spared neurons, such as CST axons that form new connections post-lesion (56, 57). Notably, Nakamura et al. showed that collateral sprouting from spared CST axons—rather than long-distance regeneration of damaged neurons— was sufficient to restore motor control (57), echoing our demonstration that MN1 can reinnervate denervated targets to restore crawling.

Additional studies highlight that promoting plasticity, not just regeneration, may be key for functional recovery (58, 59). These studies emphasize the role of the injury environment, including target denervation and glial modulation, in supporting sprouting (60, 61). Vertebrate work shows that denervated muscle and Schwann cells transiently emit growth-permissive cues that support reinnervation (62). Brown and Holland (1979) demonstrated that partial denervation of mouse soleus muscle triggers sprouting from intact motor axons, and that this response is suppressed by direct electrical stimulation of denervated muscle—indicating that specific signals from denervated targets, not simply inactivity, drive structural plasticity (63). Keynes et al. (1983) further confirmed that structurally intact, but denervated, endplates are required for nodal sprouting (64).

These parallels suggest that neuron-intrinsic identity, injury-induced cues, and the structural and molecular context of the target together shape the capacity for adaptive circuit remodeling. Our *Drosophila* model provides a genetically tractable system to dissect these mechanisms with single-cell resolution and causal precision.

### Conclusion and Perspective

Our study reveals that injury-induced heterosynaptic sprouting in the Drosophila motor circuit is triggered by complete loss of presynaptic input, is neuron-subtype specific, and contributes to functional recovery. These findings lay a foundation for uncovering the molecular logic of plasticity and circuit repair.

Drosophila offers a powerful model for dissecting the cellular and genetic mechanisms underlying this process. Future studies can use this system to identify the intrinsic growth properties of specific neurons and the extrinsic cues that promote or restrict sprouting. These insights may guide therapeutic strategies to enhance plasticity in motor disorders such as ALS and spinal cord injury, and help test whether similar principles apply in vertebrate systems.

## Materials and Methods

### Fly Husbandry

All Crosses were reared at 25°C at 50% humidity with a 12 hr light/dark cycle. For crosses with GTACR, larvae were fed with all-trans retinol (ATR) and kept in constant darkness until they were ready to be imaged or dissected. Crosses involving RPR were at 29°C to ensure ablation efficiency. For Larval ablation using Gal80ts, Embryos were kept at 25°C and as larvae hatched, they transferred to 29°C to induce ablation. A list of stocks used can be found in Table S1.

### Fillet Dissection and Immunohistochemistry

L3 larvae were used for fillet dissection, cutting on the ventral side. Dissections were fixed in 4% PFA, blocked with equal parts 5% Goat serum and 5% Donkey serum in PBST, and stained with primary and secondary antibodies. See SI Materials and methods and Table S2 for more details.

### NMJ Imaging and Quantification

Airyscan Z-stacks were acquired using a Zeiss LSM900 Confocal with a 40x Objective and processed with 3D Airyscan processing with the automatic filter. Postsynaptic Dlg of the neuron of interest were isolated using a mask of the marked neuron and the area, bouton number and bouton size were determined using a Fiji plugin developed by the Schneck lab. Branching points were counted and separated based on colocalization with Dlg and the order of branching. See SI Materials and Methods for more details.

### Behavioral Assays

Larvae were placed in groves in a 2% agarose gel and recorded (30 frames per second) using a smartphone mounted on a Zeiss Stemi 305 Microscope as they crawled 1 cm. The number of peristalsis and frames for each peristalsis were counted to determine the efficiency and duration, respectively. For analysis of overall crawling behavior, Larvae were placed on a 2% agarose gel and recorded using a smartphone above the arena for 1 minute. The WrmTrck Plugin was utilized to determine the distance traveled and for each stage the ablatio group was normalized by the average of the control. See SI Materials and Methods for more details.

### Intact Larvae Imaging and Muscle Calcium Imaging Quantification

Larvae were immobilized between a 2% agarose gel pad and a coverslip and imaged with 40x, 20x, or 10x, depending on the size of the larvae. Z-stacks of the marked axons at L0, L1, L2, and L3 were used to count the number of branching points. Time series of larvae undergoing peristalsis were analyzed using a custom MATLAB script to determine the intensity of the GCamp signal during peristalsis. See SI Materials and Methods for more details.

### Statistical Analysis

All statistical analysis was performed using R in RStudio. Averages were plotted with SEM error bars and where applicable individual samples were added as points. Number of replicates and animals is indicated in figure legends. Kruskal Wallis and Dunn post hoc were used for all NMJ quantification and behavioral data, except the data in Figure 2 which used a t-test. To compare the L1 and L3 behavioral deficits, a t-test was used. Significance was established with * = p<0.05, ** = p<0.01, *** = p<0.001, ****=p<0.0001.

## Supporting information

Detialed methods

## Acknowledgments

We are grateful to the Bloomington Drosophila Stock Center, Chris Doe, Troy Littleton, Dion Dickman, Keiko Hirono, Sarah Ackerman for generously providing transgenic fly lines. We also thank Lauren Carlisle and Elaina Hildner for their invaluable technical support. This research was funded by a grant from Texas A&M University.

---

**Fig. S1.**
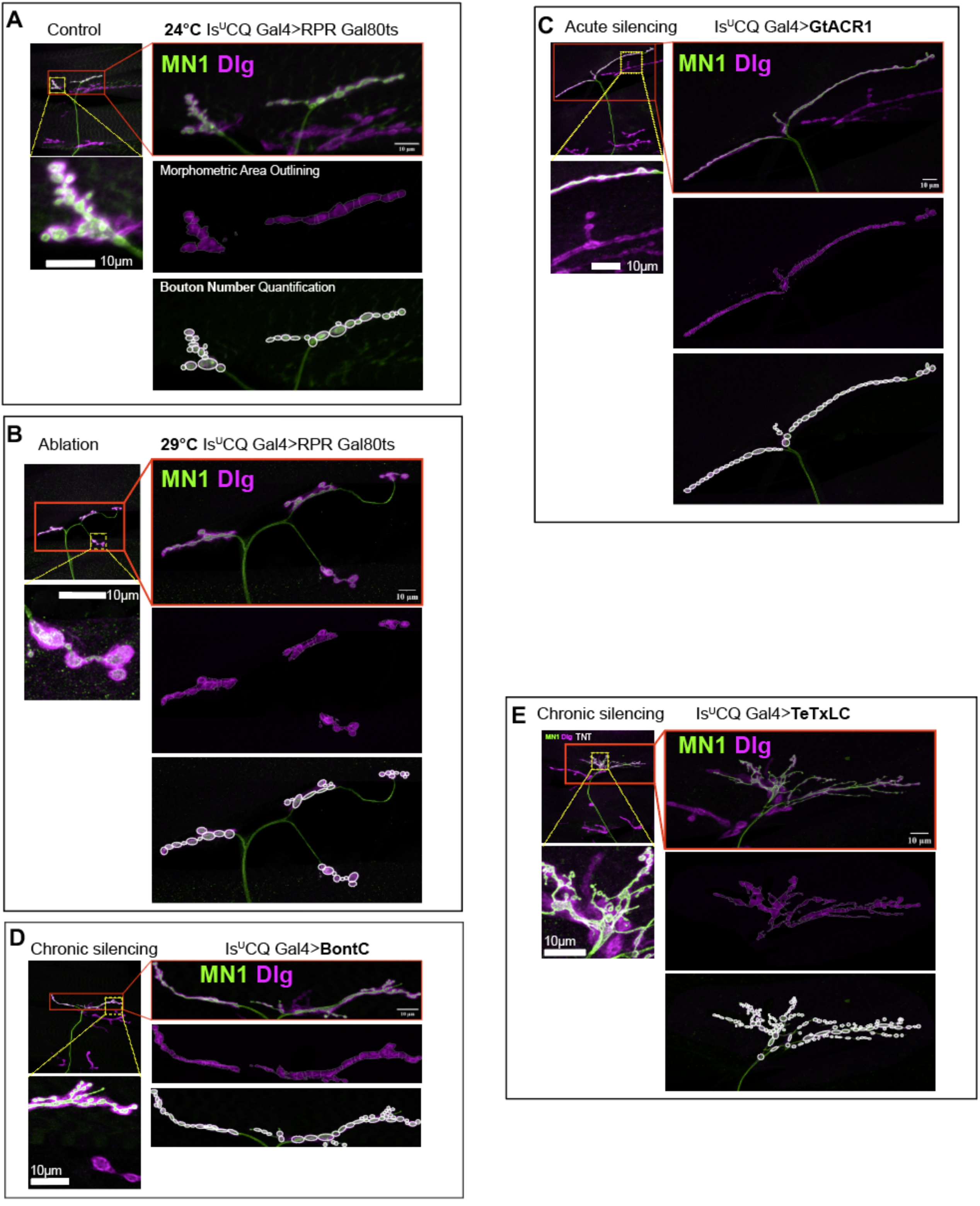
Workflow illustrating the NMJ analysis pipeline [adapted from. (**41, 42**)**] in different larval groups. Morphological responses of MN1 to distinct manipulations of neighboring motor neurons.** Representative images illustrate the structural remodeling of MN1 following ablation or silencing of adjacent tonic and phasic motor neurons (MNs), as visualized with membrane-targeted GFP and Dlg immunostaining. **(A)** In temperature-controlled **unmanipulated controls** (CQ ∪ Is-Gal4 > UAS-RPR, Gal80ts; raised entirely at 24°C), MN1 displays its typical morphology, forming a compact NMJ exclusively on its native target, Muscle 1. No ectopic projections or abnormal branching are detected. **(B)** In the **ablation group** (CQ ∪ Is-Gal4 > UAS-RPR, Gal80ts; raised at 29°C), selective apoptosis of neighboring MNs induces a dramatic response in MN1. The axon extends beyond its native territory and establishes new, ectopic NMJs on denervated Muscles 2, 9, and 10, demonstrating robust sprouting and target reinnervation. **(C)** In the **acute silencing condition** (CQ ∪ Is-Gal4 > UAS-GtACR1; light-exposed), MN1 remains restricted to Muscle 1. Although NMJ area and bouton number are elevated relative to controls, MN1 does not form branches onto adjacent muscles. **(D)** In the **chronic silencing group targeting both evoked and spontaneous release** (CQ ∪ Is-Gal4 > UAS-BoNT-C), MN1 similarly exhibits increased bouton numbers but no ectopic sprouting. The NMJ remains confined to its native muscle target, and axon branching remains limited. **(E)** In the **chronic silencing group that blocks only evoked transmission** (CQ∪Is- Gal4 > UAS-TeTxLC), MN1 shows the most extensive remodeling without sprouting. NMJ area, bouton number, and terminal branching complexity are all significantly increased compared to controls and other silencing groups. However, bouton size is notably reduced. Importantly, the tonic and phasic MNs targeted by TeTxLC appear morphologically normal (white arrows), suggesting that structural changes in MN1 are not due to degeneration of neighboring neurons but rather reflect a plastic response to altered circuit activity. Scale bars and genotype-specific labeling are provided in each panel. This figure complements the quantitative analysis shown in **main Figure 3**.

