## Supplementary material for "A Drosophila Tonic Motor Neuron Reinnervates Ectopic Muscles Fully Deprived of Native Tonic and Phasic Inputs": Detialed methods

**This PDF file includes:**

Materials and Methods
Figures S1
Tables S1 to S2
Legends for Movies S1 to S7
SI References

**Other supporting materials for this manuscript include the following:**

Movies S1 to S7

### Material and Methods

#### Fly Husbandry

All Crosses were reared at 25°C at 50% humidity with a 12 hr light/dark cycle. For crosses with GTACR, larvae were fed with all-trans retinol (ATR) and kept in constant darkness until they were ready to be imaged or dissected. Crosses involving RPR were at 29°C to ensure ablation efficiency. For Larval ablation using Gal80ts, Embryos were kept at 25°C and as larvae hatched, they transferred to 29°C to induce ablation. A list of stocks used can be found in Table S1.

#### Fillet Dissection and Immunohistochemistry

In HL3.1 (hemolymph-like solution), late 3$^{rd}$ instar larvae were pinned at the head and the tail with the ventral side up. Spring scissors were used to cut the larvae up the ventral midline and the body wall was cleaned. Four additional pins were used to splay the body wall open and the HL3.1 was removed.

Fillet dissections were fixed with 4% paraformaldehyde (PFA) for 10 minutes. After washing with PBST, samples were blocked using equal parts 5% Goat serum and 5% Donkey serum in PBST overnight at 4°C. Samples were incubated with primary antibodies (see table S2) at room temperature for 2hrs or overnight at 4°C. After washing, secondary antibodies (see table S2) were incubated at room temperature for 2hrs or overnight at 4°C. Dissections were washed in PBST, mounted in flouromont-G on microscope slides and stores at -20°C until imaging.

#### NMJ Imaging and Quantification

Using a Zeiss LSM900 Confocal microscope equipped with a 40x objective, Airyscan images were acquired in super resolution (SR) with the Zen Blue software. Raw Z-stacks were processed in Zen blue with 3D Airyscan Processing with automatic filter. Two hemisegments (A1-A7) were imaged from each animal and 6-8 animals from each condition. An Orthogonal Projection using Zen Blue was used to produce representative images.

A mask of the marked neuron of interest was used to identify the boutons belonging to the axon and all others were removed. A Fiji plugin developed by the Schneck lab [1, 2] was used to quantify the area and bouton number. ROIs were used to separately analyze the post synapses on each muscle. Bouton number was verified with a hand count based on Dlg shape and intensity. The area and bouton number from all 4 muscles were added to get the total of each. The average bouton size for each hemisegment was determined by dividing the total area by the total bouton number.

For branching quantification, segments of axon colocalized with Dlg and independent of DLG were considered separately. The number of branching points were counted and categorized based on their degree of separation from the main stalk, with direct branches being 1$^{st}$ order, branches off a 1$^{st}$ order branch being 2$^{nd}$ order, and branches off a 2$^{nd}$ order branch being 3$^{rd}$ order. All 1$^{st}$, 2$^{nd}$, and 3$^{rd}$ order branches were added to get the total number of axon or NMJ branch points.

#### Behavioral Assays

A 2% agarose gel was prepared in a 140 x 20 mm petri dish with grooves embedded using a mold. A paper with a .5 cm grid was placed beneath the gel. Individual larvae were placed in a groove using blunt forceps. Using a smartphone mounted to a Zeiss Stemi 305 Microscope, larvae were recorded at 30 fps while crawling 1cm. This was repeated for 40 larvae for each condition. The number of peristaltic waves to travel 1 cm was manually counted, efficiency is calculated by dividing the number of peristalses by 1cm. Peristalsis duration is determined by dividing the number of frames of a single peristalsis by the frame rate.

#### Intact Larvae Imaging and Muscle Calcium Imaging Quantification

To track branching across the larval stages, L0, L1, L2 and L3 larvae were washed in distilled water. Larvae were immobilized by a coverslip pressing them into a 2% agarose gel pad on a microscope slide. Depending on the size of the larvae, the 20x, 10x, or 5x objective was used to acquire a z-stack of the body wall to visualize the marked axon. Every abdominal axon from 5 animals at each stage were imaged and branching was quantified as described in NMJ quantification for each condition except branch points were not separated between axon and NMJ.

For muscle calcium imaging, the imaging was conducted with L3 larvae as described above except with a 5x objective acquiring a time series as larvae complete peristalsis. Using a custom MATLAB script [3], a ROI was placed on each muscle and placement was adjusted for each frame, avoiding overlapping sections, to track the intensity of GCamp signal during peristalsis. The calcium signal ($\Delta F/F$) for each muscle was calculated using the formula $(F - F_0) / F_0$, where F is the fluorescence at a given time, and $F_0$ is the baseline, defined as the 10th percentile of values from the first third of the recording.

**Statistical Analysis**
All statistical analysis was performed using R in RStudio. Averages were plotted with SEM error bars and where applicable individual samples were added as points. Number of replicates and animals is indicated in figure legends. Kruskal Wallis and Dunn post hoc were used for all NMJ quantification and behavioral data. Significance was established with * = $p<0.05$, ** = $p<0.01$, *** = $p<0.001$, ****=$p<0.0001$.

**Figure preparation**

Images in figures were prepared as 3D projections in FIJI (ImageJ 1.54g) and assembled using Adobe Illustrator or Adobe Photoshop.

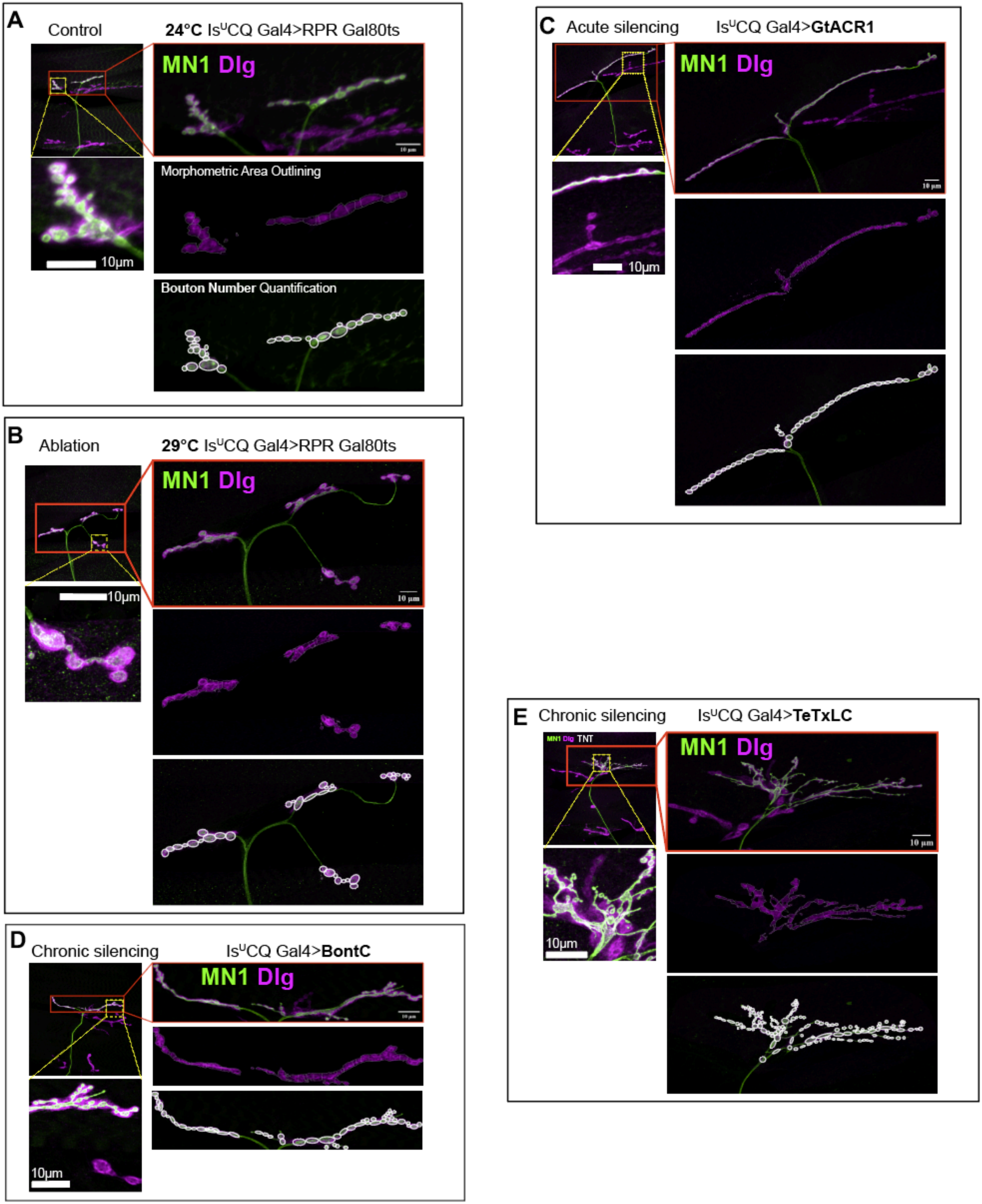

A
Control
24°C Is^U CQ Gal4>RPR Gal80ts
MN1 Dlg
10μm
Morphometric Area Outlining
Bouton Number Quantification
B
Ablation
29°C Is^U CQ Gal4>RPR Gal80ts
MN1 Dlg
10μm
10 μm
C
Acute silencing
Is^U CQ Gal4>GtACR1
MN1 Dlg
10μm
10 μm
D
Chronic silencing
Is^U CQ Gal4>BontC
MN1 Dlg
10μm
E
Chronic silencing
Is^U CQ Gal4>TeTxLC
MN1 Dlg TNT
MN1 Dlg
10μm
10 μm

**Fig. S1. Workflow illustrating the NMJ analysis pipeline [adapted from [1, 4]] in different larval groups. Morphological responses of MN1 to distinct manipulations of neighboring motor neurons.**

Representative images illustrate the structural remodeling of MN1 following ablation or silencing of adjacent tonic and phasic motor neurons (MNs), as visualized with membrane-targeted GFP and Dlg immunostaining.

**(A)** In temperature-controlled **unmanipulated controls** (CQ ∪ Is-Gal4 > UAS-RPR, Gal80ts; raised entirely at 24°C), MN1 displays its typical morphology, forming a compact NMJ exclusively on its native target, Muscle 1. No ectopic projections or abnormal branching are detected.

**(B)** In the **ablation group** (CQ ∪ Is-Gal4 > UAS-RPR, Gal80ts; raised at 29°C), selective apoptosis of neighboring MNs induces a dramatic response in MN1. The axon extends beyond its native territory and establishes new, ectopic NMJs on denervated Muscles 2, 9, and 10, demonstrating robust sprouting and target reinnervation.

**(C)** In the **acute silencing condition** (CQ ∪ Is-Gal4 > UAS-GtACR1; light-exposed), MN1 remains restricted to Muscle 1. Although NMJ area and bouton number are elevated relative to controls, MN1 does not form branches onto adjacent muscles.

**(D)** In the **chronic silencing group targeting both evoked and spontaneous release** (CQ ∪ Is-Gal4 > UAS-BoNT-C), MN1 similarly exhibits increased bouton numbers but no ectopic sprouting. The NMJ remains confined to its native muscle target, and axon branching remains limited.

**(E)** In the **chronic silencing group that blocks only evoked transmission** (CQ∪Is-Gal4 > UAS-TeTxLC), MN1 shows the most extensive remodeling without sprouting. NMJ area, bouton number, and terminal branching complexity are all significantly increased compared to controls and other silencing groups. However, bouton size is notably reduced. Importantly, the tonic and phasic MNs targeted by TeTxLC appear morphologically normal (white arrows), suggesting that structural changes in MN1 are not due to degeneration of neighboring neurons but rather reflect a plastic response to altered circuit activity.

Scale bars and genotype-specific labeling are provided in each panel. This figure complements the quantitative analysis shown in **main Figure 3**.

**Table S1.** Genetics table.

| Genetic reagent (*Drosophila melanogaster*) | Source | Description |
|---|---|---|
| *w; ;*94G06-Gal4 | Bloomington BDSC #40701[5] | 94G06 is an enhancer targeting Ib MN1 or aCC. This line expresses Gal4 in Ib MN1. Chromosome 3 |
| *w; ;*94G06-LexA | Troy Littleton [5, 6] | This line expresses LexA in Ib MN1. Chromosome 3 |
| *w;* CQ-LexA | [3, 7-13] | CQ is an enhancer that target Ib MNs (2, 3, 4, 9, and 10). This line expresses LexA in CQ MNs. |
| *w;* CQ-LexA; 94G06-Gal4 | This paper | |
| *w;* CQ-Gal4 | Miki Fujoka, Jim Jaynes BDSC #7468 | This line expresses Gal4 in CQ MNs. |
| *w; ;*27E09-Gal4 | BDSC #49227. [5] | This line expresses Gal4 in Is MNs. Chromosome 3 |
| *w;* 27E09-Gal4 | This paper | This line expresses Gal4 in Is MNs. Chromosome 2 |
| *w;* 27E09-LexA | This paper | 27E09 is an enhancer that target Is MNs (RP2 and RP5). This line expresses LexA in Is MNs |
| *w;* 27E09-LexA; 94G06-Gal4 | This paper | |
| *w;* CQ-Gal4, 27E09-Gal4 | This paper | Recombination on Chromosome 2 |
| *w;* CQ-Gal4, 27E09-Gal4; 94G06-LexA | This paper | |
| *w;* CQ-Gal4; 94G06-Gal4 | This paper | |
| *w;* CQ-LexA, 27E09-LexA | This paper | Recombination on Chromosome 2 |
| *w;* CQ-LexA, 27E09-LexA; 94G06-LexA | This paper | |
| *w;* CQ-LexA, 27E09-LexA; 94G06-Gal4 | This paper | |
| *w; ;*27E09-GFP | This paper | Expresses myr-GFP directly under the control of 27E09 enhancer. |
| *w;*13xlexAop2rpr.H}attP5/CyO; Dr/TM6C, Sb | Robin Harris, BDSC #605696 | This stock expresses apoptotic factors RPR and hid under control of LexA. |

| | | |
|---|---|---|
| w; LexAop-GtACR1.EGFP | Bloomington:92985 or 605692 | This stock expresses light gated anion channel GTACR tagged with EGFP under the control of LexA. |
| UAS-RPR.hid | [14] | This stock expresses apoptotic factors RPR and hid under control of Gal4. |
| *w;;* UAS-GTACR1-GFP | BDSC #: 92983 | This stock expresses light gated anion channel GTACR tagged with EGFP under the control of Gal4. |
| *w;* UAS-TNT.G2 | BDSC #28838 | This stock expresses the light chain of tetanus toxin (TeTxLC ) under control of UAS. |
| *w;;* UAS- BoNT-C | Dion Dickman [15] | This stock expresses botulinum neurotoxin-C (BoNT-C) under the control of UAS |
| *w;* UAS-RPR.hid, Tub-Gal80ts | Sarah Ackerman | Recombination on Chr 2 |
| *w;* UAS-RPR.hid, Tub-Gal80ts; 27E09-GFP | This paper | |
| 10×UAS-myr::GFP BDSC 32198 | Sarah Ackerman BDSC #32198 | This stock expresses the fluorescent marker GFP tagged to the membrane under the control of Gal4. |
| *w;;* LexAop-myrGFP | BDSC #32212 BDSC #32209 | This stock expresses the fluorescent marker GFP tagged to the membrane under the control of LexA. |
| *w;;* LexAop-TdTomato | BDSC # 77139 | This stock expresses the fluorescent marker TdTom under the control of LexA. |
| *w; ;*44H10::GCamp6f | [10, 16] | 44H10 is an enhancer that target muscles. This line expresses Gcamp6f in all larval muscles |
| *w;* UAS-RPR.hid,Tub-Gal80ts*;* 44H10::GCamp6f | This paper | |
| *w;;* 44H10::GCaMPf,UAS-GtACR1 | [16] | Optogenetic MN silencing along with muscle calcium imaging in crawling intact larvae. Figure 4. |
| *w;* lexAop2rpr.H}attP5*;* 44H10::GCaMPf,UAS-GtACR1 | This paper | |
| *w;* lexAop2rpr.H}attP5/CyO; 44H10::GCamp6f /TM6C, Sb | This paper | |
| *w;*13xlexAop2rpr.H}attP5/CyO; UAS-myrGFP | | |

| | |
|---|---|
| *w;* LexAop-GtACR1.EGFP /CyO; 44H10::GCamp6f /TM6C, Sb | This paper |
| *w;* CQ-Gal4,27E09-Gal4; 94G06-lexA, LexAop-TdTomato | This paper |
| *w;* CQ-Gal4, 27E09-Gal4;44H10::GCamp6f | This paper |
| *w;* 20XUAS-6XmCherry-HA}VK00018/CyO; Dr/TM6C, Sb,Tb | BDSC# 52267 |
| *w;* SP/CYO; 13XLexAop2-6XmCherry-HA}attP2 | BDSC #52271 |
| *w;* lexAop2rpr.H}attP5/CYO; 13XLexAop2-6XmCherry-HA}attP2 | This paper |

**Table S2.** Staining reagents Table

| Antibody | Host | Fluorophore | Dilution Factor | Source |
|---|---|---|---|---|
| Rabbit anti-GFP (Primary antibody) | Rabbit | n/a | 1:500 | Invitrogen |
| Mouse anti-Dlg (Primary antibody) | Mouse | n/a | 1:100 | DSHB |
| Rabbit anti mCherry/tdTomato (Primary antibody) | Rabbit | n/a | 1:500 | Novus |
| anti-F-actin (muscle) | n/a | Phalloidin CF647 | 1:40 | Biotium |
| Alexa Fluor 488 Goat anti-Rabbit (Secondary antibody) | Goat | AlexaFlour488 | 1:200 | Invitrogen |
| Alexa Fluor 555 Goat anti-Mouse (Secondary antibody) | Goat | AlexaFlour555 | 1:200 | Invitrogen |
| Alexa Fluor 555 Goat anti-Rabbit (Secondary antibody) | Goat | AlexaFlour555 | 1:200 | Invitrogen |
| Alexa Fluor 488 Goat anti-Mouse (Secondary antibody) | Goat | AlexaFlour488 | 1:200 | Invitrogen |
| Alexa Fluor 405 Goat anti-Mouse (Secondary antibody) | Goat | AlexaFlour405 | 1:200 | Invitrogen |

**Video S1. Control larva exhibits coordinated DL muscle activity during crawling.**
In an unperturbed animal (MN1 ∪ CQ ∪ Is-lexA > Aop-GtACR1, ATR–), GCaMP6f fluorescence reveals robust and sequential activation of dorsal longitudinal (DL) muscles. Calcium signals initiate in posterior segments and propagate anteriorly, reflecting normal peristaltic waves during forward locomotion.

**Video S2. Complete ablation of DL-innervating MNs abolishes DL muscle activity.**
In larvae with all tonic and phasic DL-innervating MNs ablated (MN1 ∪ CQ ∪ Is-lexA > Aop-RPR), DL muscle contractions and GCaMP6f signals are completely absent, confirming effective denervation. Activity in ventral and lateral muscles remains intact, highlighting lexA-driver specificity.

**Video S3. Acute silencing of all DL-innervating MNs eliminates DL muscle activity.**
Following light-dependent activation of GtACR1 in MN1, CQ, and Is neurons (MN1 ∪ CQ ∪ Is-lexA > Aop-GtACR1, ATR+), DL muscle calcium activity is entirely suppressed. This confirms the specificity of MN-lexA drivers and effectiveness of GtACR1 in silencing the MNs of interest.

**Video S4. Only Muscle 1 remains active when MN1 is spared from silencing.**
In CQ ∪ Is-lexA > Aop-GtACR1, ATR+ larvae (MN1 spared), calcium activity is preserved exclusively in Muscle 1, MN1's native target. DL muscles innervated by silenced MNs remain inactive, providing functional validation of MN1's exclusive input to its native muscle-1.

**Video S5. MN1-driven ectopic NMJs restore DL muscle activity following ablation.**
In CQ ∪ Is-lexA > Aop-RPR animals, where all native DL-innervating MNs except MN1 are ablated, GCaMP6f imaging reveals restored activity across Muscles 1, 2, 9, and 10. These data indicate that sprouted MN1 terminals form functional ectopic NMJs capable of driving muscle contractions.

**Video S6. Silencing the sprouted MN1 abolishes reinnervated muscle activity.**
In larvae expressing GtACR1 in MN1 after CQ∪Is-lexA > Aop-RPR-mediated ablation (CQ ∪ Is-lexA > Aop-RPR; MN1-Gal4 > UAS-GtACR1), light activation of GtACR1 eliminates all DL muscle activity. This confirms that reinnervation and functional output in previously denervated muscles depend on the activity of the sprouted MN1 terminals.

**Video S7. Peristalsis efficiency assay comparing different genotypes.**
Side-by-side recordings of larval crawling behavior under different experimental conditions illustrate the impact of MN ablation, silencing, and MN1 reinnervation on locomotor efficiency. Genotypes (left to right):

1. MN1 ∪ CQ ∪ Is-lexA > Aop-GtACR1, ATR– (control)
2. CQ ∪ Is-lexA > Aop-RPR (sprouted MN1 present)
3. CQ ∪ Is-lexA > Aop-GtACR1, ATR+ (MN1 intact but neighboring MNs silenced)
4. MN1 ∪ CQ ∪ Is-lexA > Aop-RPR (full ablation of all DL-innervating MNs)
5. MN1 ∪ CQ ∪ Is-lexA > Aop-GtACR1, ATR+ (all DL MNs acutely silenced)
6. CQ ∪ Is-lexA > Aop-RPR; MN1-Gal4 > UAS-GtACR1, ATR+ (sprouted MN1 acutely silenced)

This assay visually demonstrates the functional consequences of MN ablation, silencing, and reinnervation on coordinated locomotor output.
